# Development and application of a genotyping by target sequencing SNP array panel in *Salix suchowensis*

**DOI:** 10.1101/2025.05.12.653369

**Authors:** Yu Han, Shaobo Gu, Mingjia Zhu, Wei Liu, Landi Feng, Tongming Yin, Xuemeng Gao, Yanjun Zan, Yan Ji, Jianquan Liu

## Abstract

*Salix suchowensis* is an important species of *Salix*, known for its rapid growth property and wide application in environmental construction, ecological restoration, wicker production, and biomass energy production. Due to its significance as a sustainable biological resource, *S. suchowensis* has been the centre of intensive breeding. However, rapid improvement of growth and biomass has been hindered by a lack of genomic resources. To address this limitation, we designed a genotyping by target sequencing SNP array panel to facilitate efficient genotyping. Using whole-genome resequencing data, a total of 39,076 SNPs were selected for the array panel, consisting of trait-associated SNPs, intragenic SNPs, and intergenic SNPs. This panel was validated by genotyping 550 new samples, demonstrating high call rates and effective capture of the population structure. Genome-wide association analysis identified 72 SNPs associated with plant height and ground diameter. Additionally, the array panel shows a high potential for genomic selection, with high prediction accuracy for various traits. These results highlight the efficiency of this panel in capturing genomic variations that are highly valuable for future genetic research and breeding applications.

## Introduction

Willows (*Salix*) are dioecious, catkin-bearing woody plants, together with poplars (*Populus*), which comprise the *Salicaceae* family. Willows consist of more than 300 species, including shrubs and trees, which are widely distributed in diverse ecological niches, particularly in temperate and arctic zones. Asia is considered the centre of origin for *Salix*, with China alone harbouring around 270 species (Newsholme, 2003; Argus, 1997). The physiological adaptations and ecological resilience of willows make them ideal candidates for ecological conservation and restoration projects across various climatic zones. Moreover, willows have a significant commercial impact, especially in wood production. Their rapid growth and short generation period make them an ideal renewable biological resource for biomass production, with additional economic value in manufacturing, medicinal use, and ornamental cultivation (Freudl, 2018; Kuzovkina and Quigley, 2005; Christersson *et al*., 1993; Elowson, 1999). Although willows have a long breeding history, genetic improvement has been limited by a lack of genomic resources and a poor understanding of the genetic basis of growth traits. Recently, genomics-informed breeding strategies have been widely applied to a variety of crops (Varshney *et al*., 2021; Wang *et al*., 2025; Alemu *et al*., 2024; Basu and Parida, 2021). Adopting this approach in willow breeding could facilitate rapid genetic improvement of key growth traits.

*Salix suchowensis* is the first *Salix* species with a high-quality reference genome, which has annotated more than 30,000 protein-coding genes (Dai *et al*., 2014; Wei *et al*., 2020). This makes it an ideal model for functional genomics and genetic studies in willows. Growth traits are key agronomic characteristics that directly influence biomass production and are primary targets for genetic improvement. Previous studies have identified quantitative trait loci (QTLs) underlying growth traits in *S. suchowensis* using amplified fragment length polymorphism markers and genotype by sequencing (Freudl, 2018; Freudl, 2018). However, there is an urgent need for a high-throughput, cost-effective, time-saving, and accurate genotyping method for large-scale population research in *S. suchowensis*. Targeting by sequencing is a promising method to provide high-resolution single-nucleotide polymorphisms (SNPs), which can be used for genome-wide association studies (GWAS), genomic selection(GS), and population genetic analysis (Thomson, 2014; Balagué-Dobón *et al*., 2022; Kim *et al*., 2022; Freudl, 2018; Johnson *et al*., 2018). The up-to-date technology is genotyping by target sequencing (GBTS), which enhances flexibility and scalability since the designed probes can be easily updated and expanded (Freudl, 2018; Freudl, 2018). GBTS SNP panel has been successfully applied in various species, including rice, maize, and soybean, proving its efficiency in genetic research (Freudl, 2018; Freudl, 2018; Guo *et al*., 2021; Lee *et al*., 2022; Freudl, 2018). Thus, the development of a GBTS SNP panel for *S. suchowensis* would be a valuable resource for genetic research and breeding programs.

In this research, we designed a 40k GBTS SNP panel for *S. suchowensis* based on whole-genome resequencing data. This SNP panel was designed to capture the genetic diversity for genetic research and breeding applications. We validated this SNP panel by genotyping 550 new samples and conducting GWAS to identify candidate genes associated with growth traits. Finally, we assessed the prediction accuracy of the SNP panel for various complex traits in *S. suchowensis*.

## Materials and Methods

### Selection of 40K SNPs from WGS

A total of 4,531,211 SNPs obtained from whole-genome resequencing in our previous study were used. All SNPs are bi-allelic variants with a call rate above 90% and a minor allele frequency (MAF) greater than 0.05. In 2023, 600 samples from 8 families were planted in Leshan (LS; N29.55 °, E103.77°), Pengzhou (PZ; N30.99°, E103.96°), and Yibin (YB; N28.75°, E104.64°). Phenotypic data for plant height (PH) and ground diameter (GD) were collected from all three locations, and transcriptome data were gathered in LS to perform GWAS and expression quantitative trait loci (eQTL) mapping. In total, 25 SNPs associated with growth traits and 938 SNPs associated with gene expression were prioritised in the SNP panel design.

Moreover, we categorised these SNPs based on their relative genomic position with gene annotation. Considering the potential significant influence of SNPs located at the start and stop codons, such SNPs were included in the SNP panel design (239 at the start codon and 82 at the stop codon).

Then, 3,235,172 SNPs located within gene regions (including gene body, 3 kb flanking region of upstream and downstream) were further filtered to ensure at least one SNP per gene, thereby maximising the coverage of regions with annotated genes. For each gene, SNPs inside the gene region were used to calculate pairwise R^2^ values. After that, SNPs with the top 10 largest number of tagged SNPs (R2>0.6) were retained. The threshold we selected under the consideration that these SNPs could best represent the genetic variations inside corresponding genes. Next, the SNP with the highest MAF was chosen for each gene, for its high occurrence rate in this population.

Furthermore, for the 1,296,039 SNPs located outside the gene region, linkage disequilibrium (LD) filtering was performed to obtain representative SNPs. Using PLINK with a 200 kb window, a 10 kb stepsize, and an R^2^ threshold of >0.1, we identified 124,192 SNPs. These SNPs were subsequently down-sampled to 12,419 SNPs (Chen *et al*., 2019).

Based on the preliminary selection of SNPs, we designed the liquid-phase probe sequences located within the upstream and downstream regions of the SNP loci. The probe sequences were designed using Novogene’s SNP array algorithm based on thermodynamic stability principles to ensure high capture efficiency. The probe length was set between 80-120 bp with an average of approximately 100 bp, while maintaining GC content within 20%-70%. The number of SNPs in probe capture regions was restricted to fewer than 5. The SNPs of the panel that do not meet the standards will be filtered. This step ensured high capture efficiency of this SNP panel. Finally, 39,076 SNPs—nearly 40k—were retained as the final SNP panel.

### Plant materials

After designing the 40k SNP panel, we randomly selected two samples from each of the previously research samples, resulting in a total of 16 samples, to validate the quality of the SNP panel. These samples were collected from an experimental field in 2023, which contained 600 samples. The youngest, most recent leaves from the tips of the main stems were collected and immediately frozen in liquid nitrogen for DNA extraction.

In this study, 550 samples from 8 new full-sib families of *S. suchowensis* were selected to perform GBTS and genetic investigation. These samples were planted at two locations, Yibin (YB; N28.75°, E104.64°) and Pengzhou (PZ; N30.99°, E103.96°) cities, in Sichuan province, China. Fifteen days after planting, two to three fresh leaves were collected from each sample to serve as material for DNA extraction. The collected leaves were immediately frozen in liquid nitrogen to preserve DNA integrity. Phenotypic data collection began 30 days after planting, at a point when plants exhibit measurable differences in growth traits. At this stage, we take measurements of plant height (PH) and ground diameter (GD) at 15-day intervals at both YB and PZ locations until the end of the growing season. PH was measured as the vertical distance from the soil surface to the tip of the main stem (apical meristem). GD was determined as the diameter of the stem at approximately 5 cm above the soil surface. These two traits measured the vertical and horizontal growth of the plant, capturing major traits related to biomass accumulation.

### Validation of the 40k SNP panel

All samples were sequenced using GBTS, with an average of 2 Gb raw reads per sample. The raw data were processed using Fastp to filter out low-quality reads and adapters (Chen, 2023). Clean data were then aligned to the reference genome of *S. suchowensis* using BWA-MEM (Li, 2013). The aligned data were processed with SAMBAMBA to sort and remove duplicate reads. Genome coverage was calculated using SAMtools (Li *et al*., 2009; Tarasov *et al*., 2015). Genetic variants were detected using BCFtools (Danecek *et al*., 2020). The genotyping data were filtered with the following parameters: QUAL > 30, F_MISSING < 0.5, DP > 2, and “-m2 -M2”. SNPs located at the designed positions were extracted for further analysis, including the calculation of the sample call rate and site call rate. The sample call rate is defined as the average SNP call rate across all SNPs for a given sample, while the site call rate refers to the SNP call rate across all samples. For further analysis, SNPs with MAF < 0.05 were excluded using PLINK (Chen *et al*., 2019).

### Principal component analysis

The population structure of the 600 *S. suchowensis* individuals was evaluated using principal component analysis (PCA) based on the whole-genomic SNPs and the 40k selected SNPs. PCA was performed using the GCTA software (Yang *et al*., 2011). For whole-genomic SNP data, we select all SNPs from the previous study as input to perform PCA analysis. For the 40k selected SNPs data, we extracted these SNPs from whole-genomic SNPs as input to perform PCA analysis. The PCA results were visualised by R language.

### Genetic analysis

The kinship heritability for each trait was estimated by fitting a mixed linear model using GCTA software (Yang *et al*., 2011).

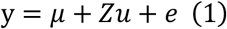

Here y is the phenotype value, *µ* is the intercept of the model. *Z* is the corresponding design matrix that satisfies *ZZ*^*T*^/_*m*_ = *G*, where *G* is the kinship matrix, *m* is the SNP number. Therefore, the distribution of *u* is 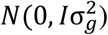 is the residual with 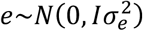. The kinship heritability was calculated as 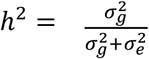. For comparison, the genomic kinship matrix G was calculated based on the whole-genome SNPs and the 40k selected SNPs seperatly. Phenotypic data were collected from previous study. After planting 550 new samples in YB and PZ, PH and GD were collected to perform a genome-wide association study (GWAS). GWAS was performed using GEMMA with a mixed linear model (Zhou and Stephens, 2012). A Bonferroni-corrected genome-wide significance threshold (0.05 divided by the number of SNPs) was used for genome-wide significance. The significantly associated SNPs were annotated using the *S. suchowensis* genome annotation file. Genes within a 10 kb flanking region upstream and downstream of the associated SNPs were extracted for gene ontology (GO) enrichment analysis using the clusterProfiler package in R (Wu *et al*., 2021).

### Genomic Selection

To assess the quality of this SNP panel in GS, the Genomic Best Linear Unbiased Prediction (GBLUP) method was employed. This method, described in model 1, integrates additive genetic effects into a mixed linear model framework, enabling the estimation of genomic estimated breeding value (GEBV) based on SNP-derived relationships. This model was implemented in the hglm package in R (Alam *et al*., 2015). The G matrix was calculated based on the 40k selected SNPs. To evaluate the accuracy of genomic selection, five-fold cross-validation was used to split the dataset into training and testing sets. For each fold, GEBVs were calculated for individuals in the testing set based on the G matrix and model parameters derived from the corresponding training set. The correlation between the calculated and predicted breeding values across the five test sets was used to assess the accuracy of genomic selection.

## Results

### Probe design of *S. suchowensis*

A total of 4,531,211 high-quality SNPs from 600 *S. suchowensis* F1 individuals were obtained for probe design. Four selections were applied to select the SNP panel (Fig 1): (1) SNPs associated with growth traits (25 SNPs) and molecular traits (938 SNPs) identified through GWAS and eQTL mapping were prioritised for their functional relevance. (2) SNPs located in the start and stop codons (239 and 82 SNPs, respectively) were included due to their potential role in gene regulation. (3) At least one SNP per gene within the gene body were selected to ensure maximum gene space coverage. (4) SNPs were evenly distributed in intergenic regions to account for the even genome distribution of SNP panel. Finally, after filtering SNPs by their phychemical properties (see materials and methods), we retained 39,076 probes for the final SNP panel (Supplementary Table 1), which were evenly distributed across the genome (Fig 2A). The number of SNPs per chromosome is highly correlated with chromosome length (Pearson correlation coefficient = 0.96, *p*-value = 1.07e-10), with the highest number of SNPs on chromosome 1 (4,401 SNPs) and the lowest number on chromosome 9 (938 SNPs) (Fig 2B). With MAF higher than 0.05 as the selection criterion, the median MAF of the 40k SNP panel was 0.43 (Fig 2C), which ensured the selected SNP panel SNPs were widespread in the population. These SNPs are located in different genomic regions, including the gene body and intergenic region (Fig 2D). Following our selection criteria, most of the selected SNPs are distributed within the gene structure, with the most SNPs located in the exon region (11,728 SNPs).

**Figure 1.**
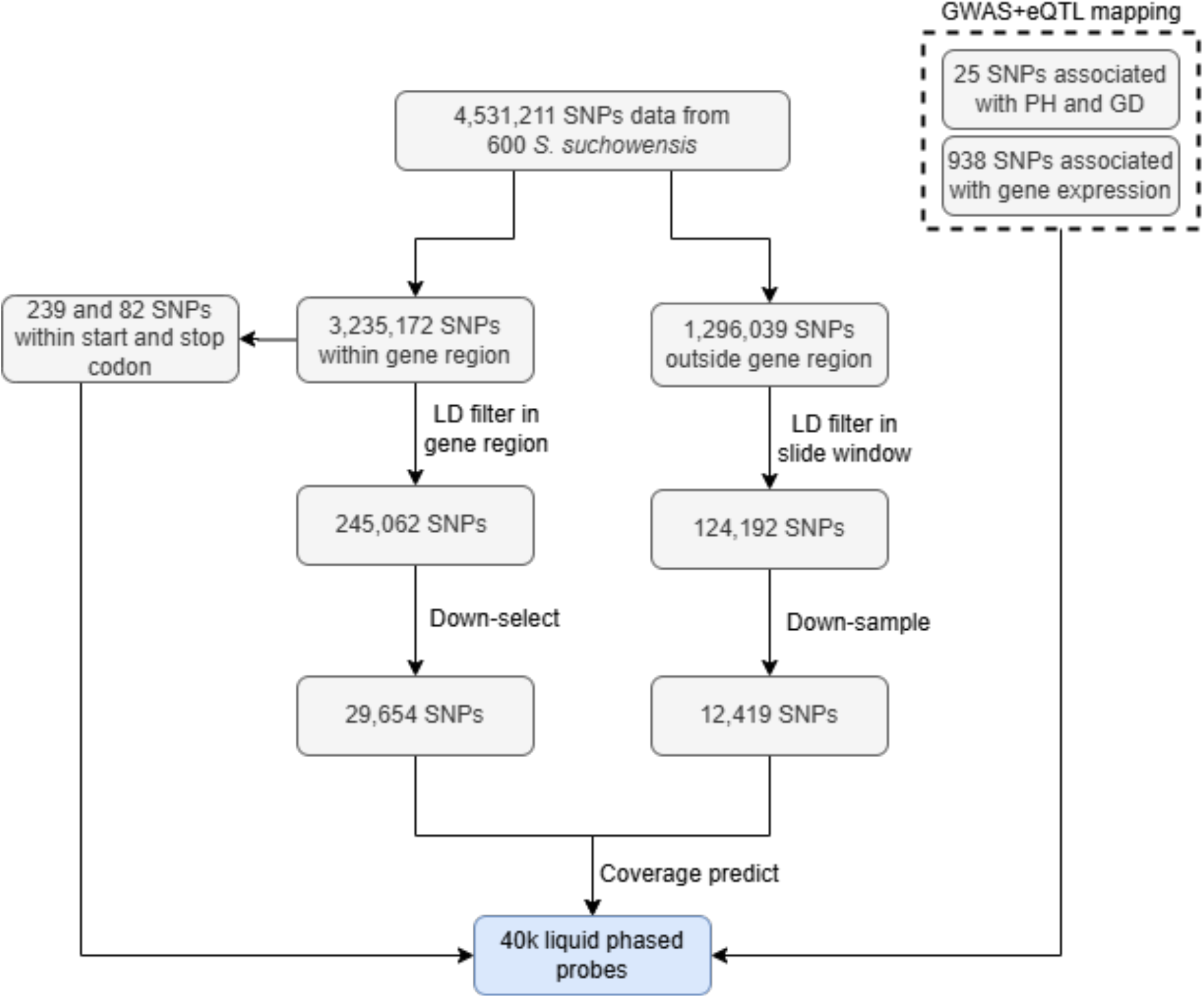
Schematic representation of the probe design workflow. Each box denotes the number of SNPs after each selection step. The abbreviated selection steps from the materials and methods are indicated around the arrows.

**Figure 2.**
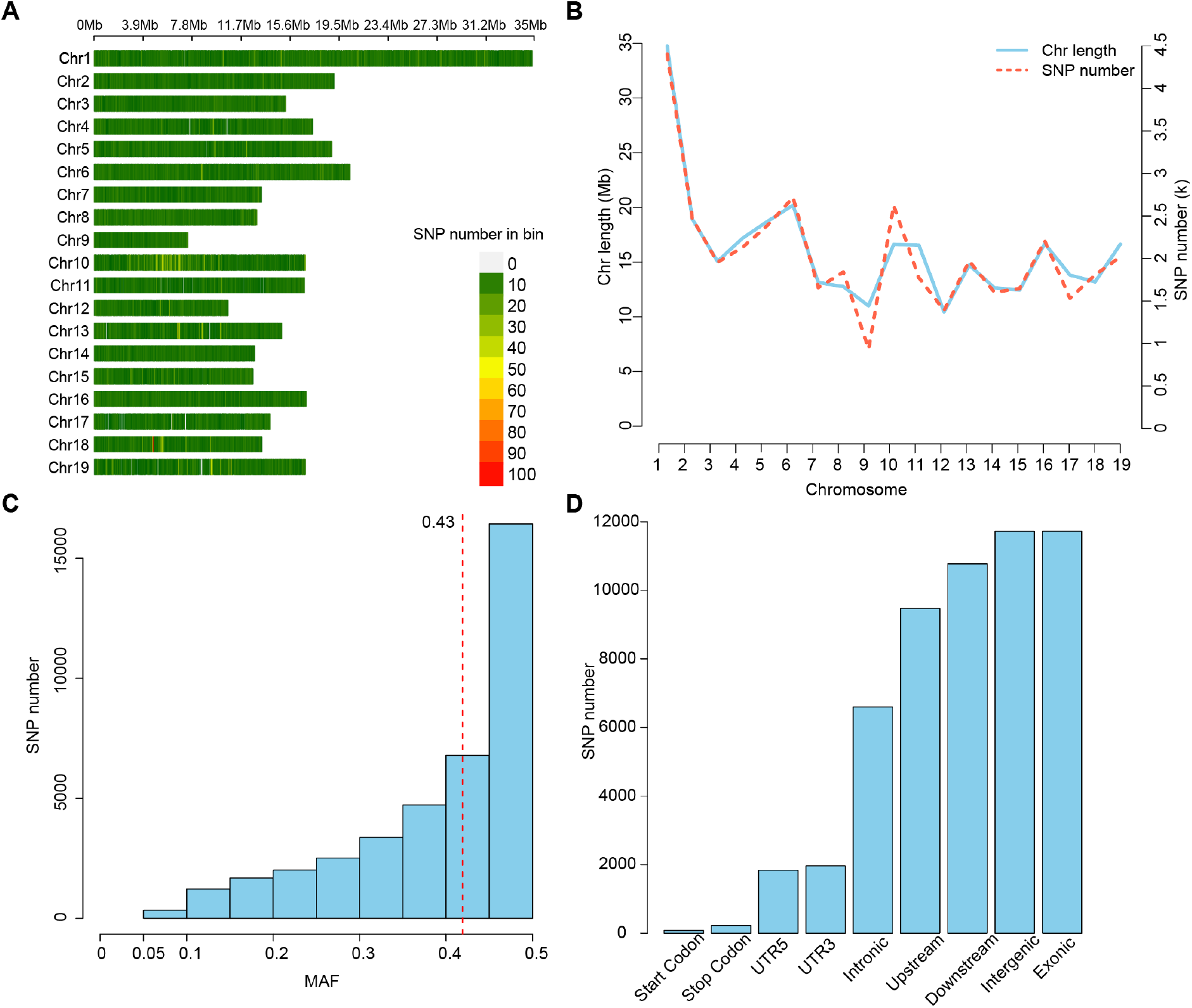
Characteristics of the 40k probes. **(A)** The distribution of SNPs by chromosome. Each bar represents a 50 kb genome bin, colored by the number of SNPs. **(B)** The plot of chromosome length (colored in blue) and the number of SNPs per chromosome (colored in red). **(C)** MAF of the SNP panel. **(D)** Distribution of SNPs with different genomic features.

### 40k SNPs captured the population structure of *S. suchowensis*

To evaluate whether the 40k selected SNPs could capture genetic relatedness and diversity of the *S. suchowensis* population, we compared the population structure of the 600 *S. suchowensis* captured by whole-genomic SNPs and the 40k selected SNPs. PCA analysis indicated a similar population structure in each family (Fig 3A, B). In the PCA plots, clear separations along the first and second principal components reveal distinct genetic signals for each family, with clusters displaying consistent internal distribution and minimal overlap. Consistent with that, the 600 individuals from eight families, individuals can be clearly separated into eight groups in the PCA plot, both by the whole-genomic SNPs and 40k selected SNPs. This suggests that the selected SNPs are highly efficient in capturing genomic relationships among the studied individuals.

**Figure 3.**
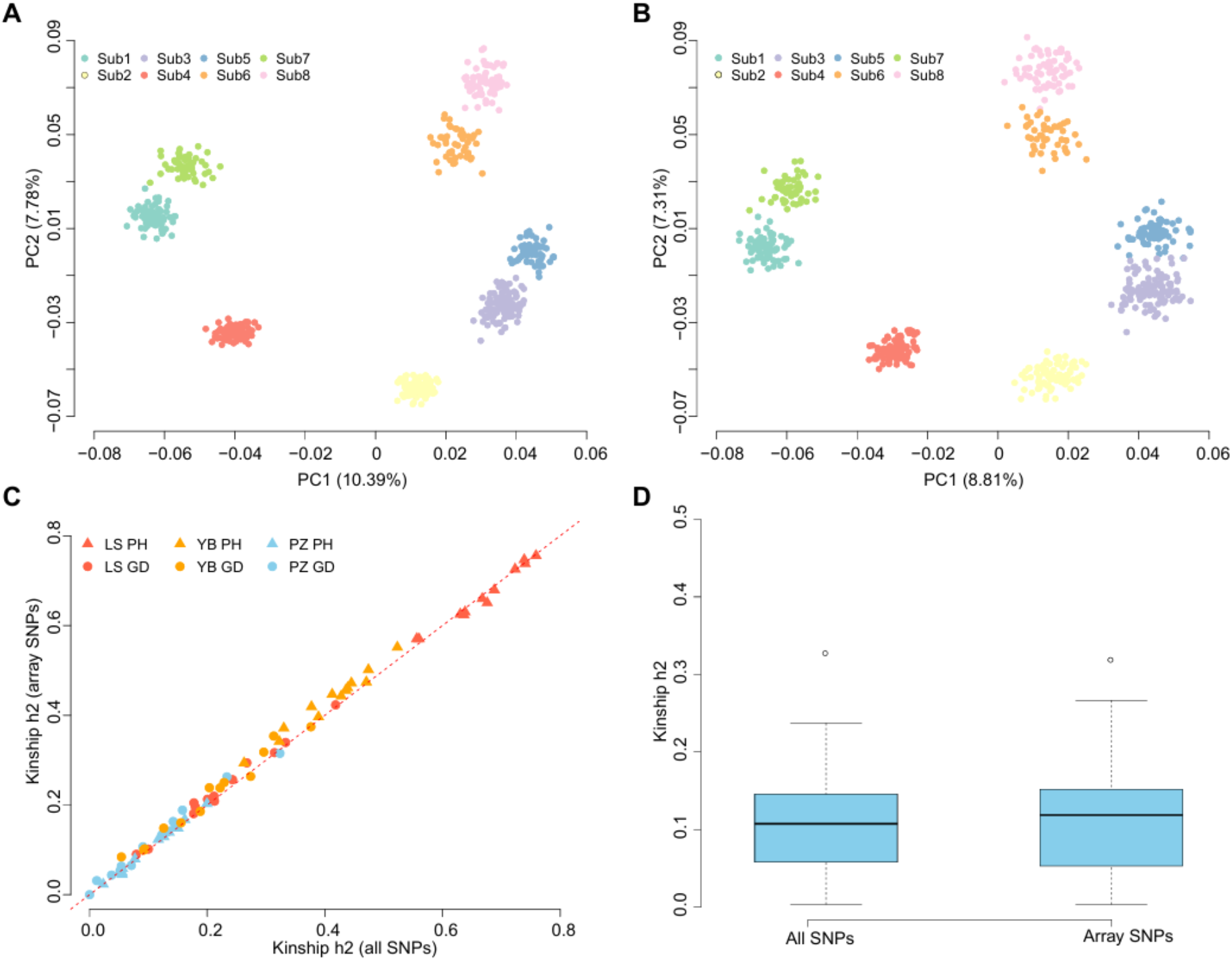
The 40k SNP panel captured the population structure of *S. suchowensis*. **(A)** and **(B)** PCA plot of the 600 *S. suchowensis* individuals based on the whole-genomic SNPs and the 40k selected SNPs. Each color represents a family. **(C)** The estimated kinship heritability of growth traits was based on the whole-genomic SNPs and 40k selected SNPs. Different color and point shapes were used to indicate the source of the phenotype data from the previous study. The red line represents the diagonal line. **(D)** The boxplot of the heritability of all traits in Pengzhou.

Furthermore, we compared the estimated kinship heritability of measured growth traits (PH and GD at YB, PZ, and LS) to assess the quality of selected SNPs in genetic research. This comparative analysis was performed across multiple environments and traits, ensuring a comprehensive evaluation under a wide range of scenarios. Similarly, the estimated heritability of growth traits from the whole-genome sequenced SNPs and 40k selected SNPs was highly correlated (Pearson correlation coefficient = 0.99, *p*-value = 2.82e-93, Fig 3C). For instance, at Pengzhou, the difference in heritability median between whole-genome sequenced SNPs and 40k selected SNPs was just 5.68e-3. Such subtle differences highlighted the high quality of the 40k SNP SNP panel in capturing the genetic variance of studied traits, making it an ideal approach for future genetic research and breeding applications.

### Validation of the probe efficiency

Here, we selected two individuals from each family to validate the capture ability of synthetic DNA probes. As illustrated in Fig 4A, the SNP panel showed a high call rate across all selected samples, with a minimal call rate of 96.95%, and the combined SNP call rate reaches 99.91%. Then we applied this SNP panel in a breeding population with 550 samples from eight additional families by performing GBTS. With ∼2 Gb raw sequencing reads per individual, 50% of samples showed an average coverage of 219.70X at targeted positions (Fig 4B). The high sequencing, with minimal fluctuations in coverage across targeted probe regions, ensured robust SNP calling for downstream analysis. For each probe, the sequence depth showed a normal distribution, with a median depth of 88.49X, and gradually decreased towards flanking region (Fig 4C). These results demonstrated that the designed probes could bind DNA with high accuracy and specificity. After SNP calling, a high call rate (median 97.05%) across all samples was achieved (Fig 4D). Moreover, for designed probes, 50% of probes showed a high call rate across all sites (median 99.82%) (Fig 4E). Altogether, these results indicated a high efficiency of the designed SNP panel.

**Figure 4.**
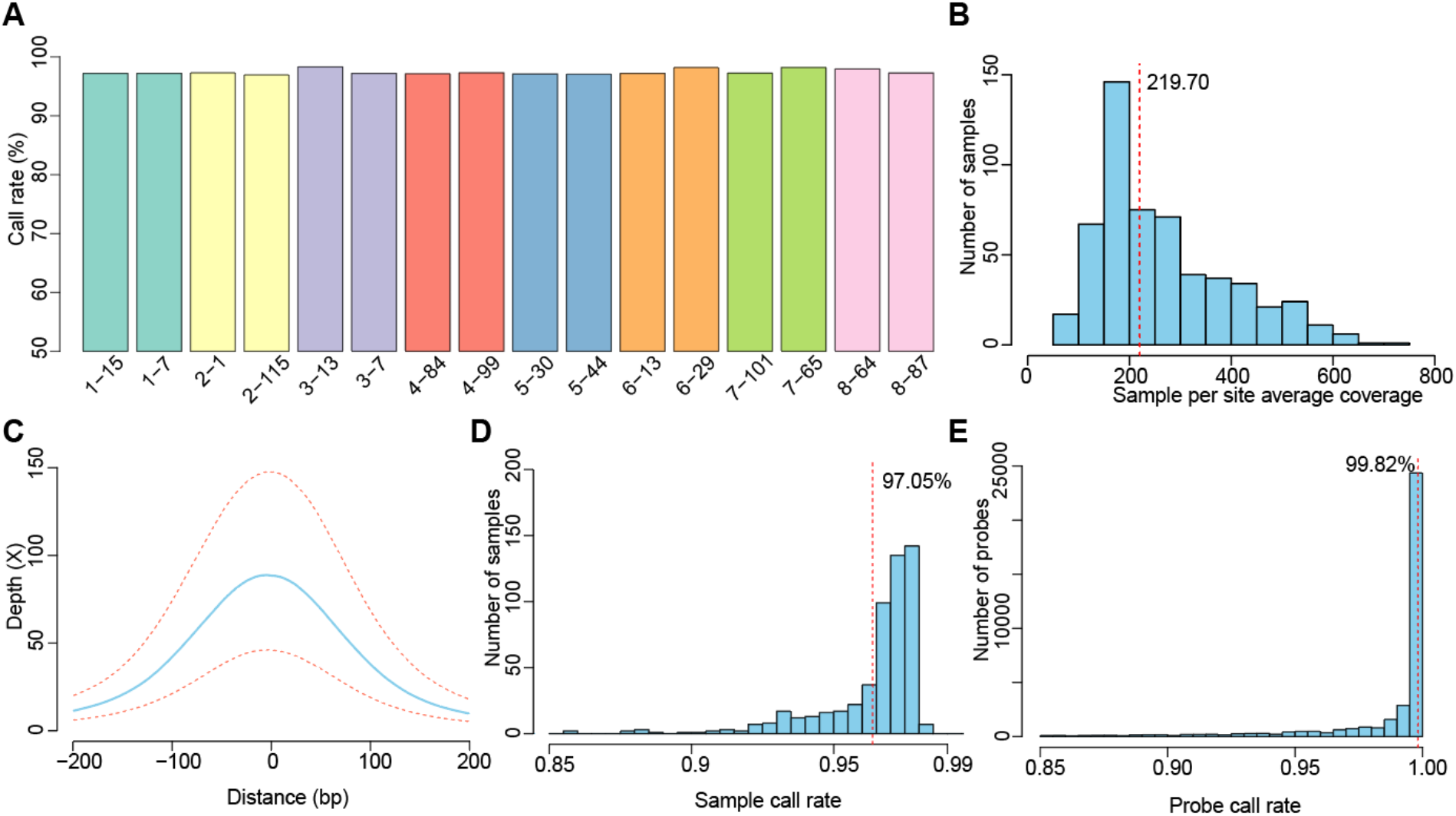
Validation of the probe efficiency. **(A)** The call rate of the 40k SNP panel in 16 selected samples. The color of each bar represents the family, which is consistent with Figure 3 **(B)** The coverage of the 40k SNP panel in 550 new samples. The red line indicates the median value. **(C)** The distribution of median sequence depth of the 40k SNP panel in 550 new samples across 400bp base (blue). Distance 0 represents the selected snp position, upstream was shown as minus, and downstream was shown as positive number. The red line above the blue line indicates the top 25% depth of each base, and the red line below the blue line indicates the bottom 25% depth of each base. **(D)** The sample call rate of the 40k SNP panel in 550 new samples. The red line indicates the median value. **(E)** The site call rate of each 40k SNP panel. The red line indicates the median value.

### Applying the SNP panel in GWAS to identify candidate genes for growth traits

To explore the implementation ability of this SNP panel in dissecting the genetic architecture of growth traits, we measured the phenotype of PH and GD on 550 samples at 15-day intervals in YB and PZ from one month after planting (Supplementary Table 2), and performed GWAS analysis. A total of 72 SNPs were significantly associated with growth traits (Fig 5A-D, Supplementary Table 3). There were 63 SNPs distributed inside the body of 50 genes, and nine SNPs were located at intergenic regions (Fig 5E). We noticed that there were three SNPs located in the stop codon, which may introduce gene expression change and further influence the growth traits. These rusults highlighted the significance of the SNP panel in dissecting the genetic basis of growth traits. For instance, SNP 4:16980574 (G→A) is located in the stop codon of *IMY05_004G0170200. IMY05_004G0170200* is annotated as a vacuolar-sorting receptor, which recognizes specific protein sorting signals.Such variation may affect plant growth by mediating the sorting and trafficking of proteins to vacuoles (Masclaux *et al*., 2005). Within 10 kb flanking region of all associated SNPs, 169 genes were detected. GO enrichment analysis indicated that these genes were enriched in the biological process of cell division, cell development, and growth (Fig 5F), suggesting they were promising candidates for future validation.

**Figure 5.**
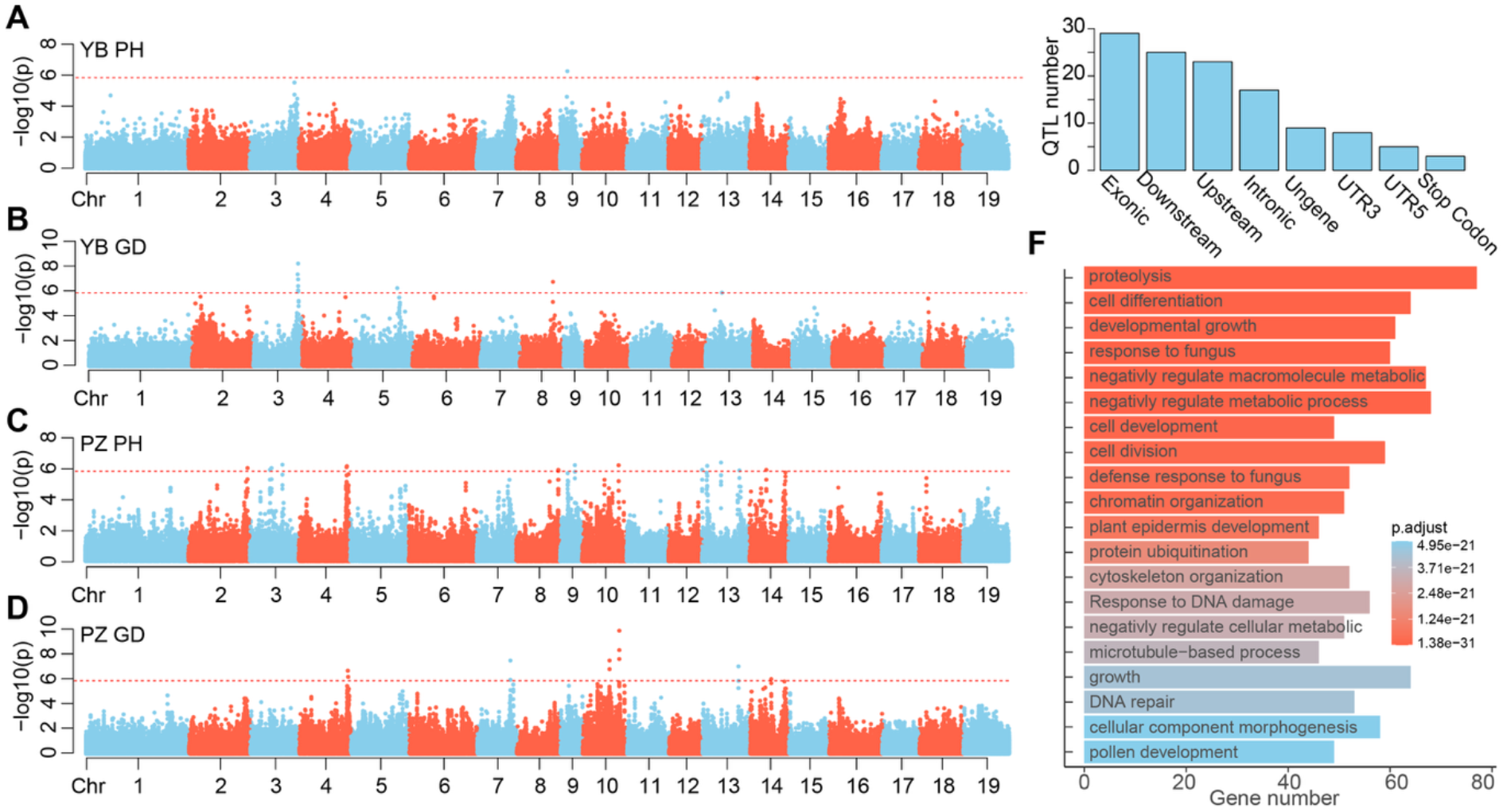
GWAS of growth traits identified candidate genes. **(A-D)** The overlapping Manhattan plot of GWAS for the five measured plant heights (PH) and ground diameter (GD) in Yibin (YB) and Pengzhou (PZ). The red line represents the Bonferroni corrected genome-wide significance threshold. **(E)** The distribution of significantly associated SNPs in gene and intergenic regions. **(F)** GO enrichment analysis of genes within 10 kb flanking region of significantly associated SNPs.

### The designed SNP panel enables accurate genomic selection in *S. suchowensis*

GS is a efficient approach in rapid genetic improvement of livestock, crop and forest trees. It could significnalty improve breeding efficiency by selecting the elite individual without measuring phenotype. After using five-fold cross-validation to split the training and testing set, the Pearson correlation of the calculated and predicted breeding value in five test sets was visulised. As showed in Fig 6, all the traits measured at different plant locations and time showed a high prediction accuracy, with a median of 0.90. This high accuracy demonstrated the high efficiency of this SNP panel in genomic selection in *S. suchowensis*, highlighting the potential of this SNP panel in future breeding applications.

**Figure 6.**
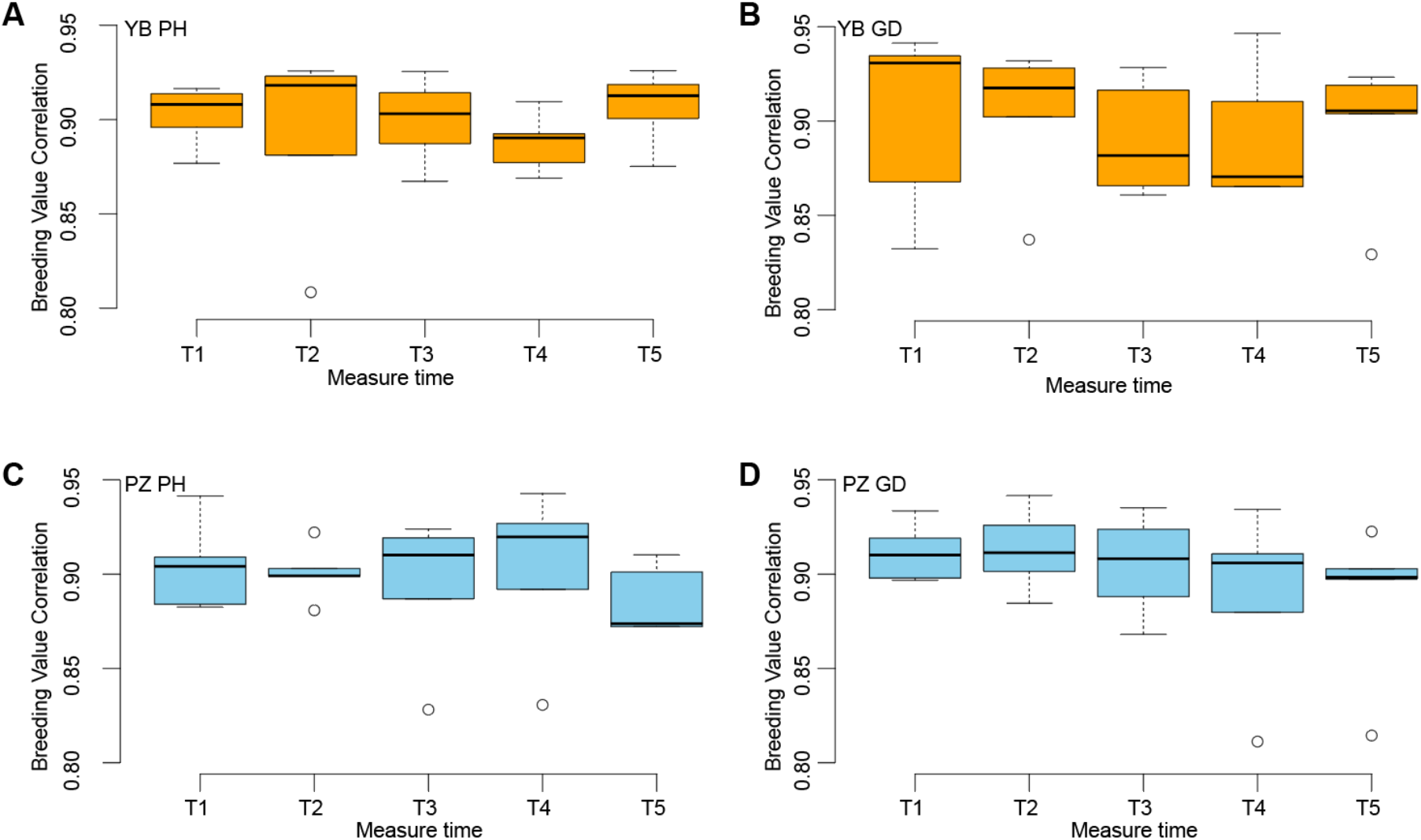
The accuracy of genomic selection in *S. suchowensis*. **(A-D)** The boxplot of correlation of calculated and predicted breeding value in five test sets for plant height (PH) and ground diameter (GD) in Yibin (YB) and Pengzhou (PZ).

## Discussion

### The design and performance of the SNP panel

Common strategy for SNP panel design typically focuses on evenly distributing SNPs across the genome, aiming at maximizing the genome coverage (Freudl, 2018; Freudl, 2018). In this study, we adopted a strategy that prioritizes the biological function of SNPs, while maintaining even distribution across the genome. As described above, these selection criteria consider candidate SNPs that may influence growth traits and gene expression, SNPs that may have effect on gene translation, the most representative SNPs located within gene body, and SNPs that fill gaps of the selected SNPs in terms of genome coverage. Ultimately, 39,076 SNPs were selected. The consistent population structure and high kinship heritability captured by both whole-genome SNPs and the SNP panel indicte a high quality of the designed SNP panel. The SNP panel’s performance was furthur validated by genotyping 550 samples of *S*.*suchowensis*, yielding a high call rate across all samples (median 97.05%) and all sites (median 99.82%). These high call rates demonstrate the efficiency of the SNP panel in genotyping. Additionally, when compared to solid-phase gene chips, the GBTS SNP panel enhances the flexibility of probe design and provides more comprehensive information (Freudl, 2018; Dalma-Weiszhausz *et al*., 2006). For instance, additional SNPs can be easily incorporated into probe designs for further population studies. Moreover, the target sequencing can capture a wide range of genomic variants in the probe region, offering many kind of molecular marker data, such as SNPs, SSRs, and InDels, and providing a much larger number of markers than that obtained from multiplex PCR (Freudl, 2018).

### Application of SNP panel in genetic research and breeding

As genetic research in willow has not progressed as rapidly as in other crops, and the first high-quality reference genome for the model species *S*.*suchowensis* was only published five years ago, the molecular mechanisms and genetic architecture of *S*.*suchowensis* remain unclear. The 40k SNP SNP panel designed in this study provides a valuable resource for advancing genetic research in this species. The SNP panel was successfully applied in GWAS to identify QTL associated with growth traits. A total of 72 SNPs significantly associated with growth traits were identified, and genes around these SNPs were enriched in biological processes such as cell division, cell development, and growth. These findings lay the foundation for unravelling the genetic architecture of growth traits in *S. suchowensis*.

The Omnigenic model is proposed for the genetic mechanism of complex traits, which describes that complex traits are not only directly affected by a few core genes from the core pathways, but also indirectly regulated by a large number of peripheral genes (Boyle *et al*., 2017; Mathieson, 2021). Core genes are usually located at the central nodes of biological networks and play a key regulatory role in phenotypes, while peripheral genes regulate the topological structure of the network by influencing or being affected by core genes. These genes collectively shape the variation of phenotype. In the GWAS analysis results, the genetic structure of the ground diameter and plant height traits embody this model in *S. suchowensis*. Although some genes near SNP loci are functionally candidate genes for regulating growth and development, there are still some SNP loci significantly related to traits distributed in intergenic regions or other functional genes. These core genes may have regulatory relationships with a large number of peripheral genes, influencing the core growth pathway through a complex gene network. However, by implementing this SNP panel, large-scale SNP genotyping can be achieved at a lower cost, thereby providing an opportunity to accelerate the genetic dissection of the growth traits.

Moreover, the high accuracy of genomic selection observed in this study (median correlation of 0.90) further demonstrates the feasibility of using this SNP panel for GS. With continuous updates of the SNP panel and increasing amount of genotype and phenotype data, prediction models can be further optimized to improve GS accuracy and accelerate genetic improvement of *S. suchowensis*.

## Conclusion

We designed a 40k SNP panel for *S. suchowensis* by including trait-associated SNPs, gene body SNPs, intragenic SNPs, and intergenic SNPs, which are evenly distributed across the genome. We performed a comprehensive evaluation analysis and revealed its high quality in capturing population structure, identifying QTL associated with growth traits and predicting complex traits. Overall, our analysis demonstrated the potential of this SNP panel in genetic studies and breeding applications, which could make a valuable contribution to a wide range of applications.

## References

Alam, M., Rönnegård, L. and Shen, X. (2015) Fitting conditional and simultaneous autoregressive spatial models in hglm. R J, 7, 5.

Alemu, A., Åstrand, J., Montesinos-López, O.A., et al. (2024) Genomic selection in plant breeding: key factors shaping two decades of progress. Mol. Plant, 17, 552–578.

Argus, G.W. (1997) Infrageneric Classification of Salix (Salicaceae) in the New World. Syst. Bot. Monogr., 52, 1–121.

Balagué-Dobón, L., Cáceres, A. and González, J.R. (2022) Fully exploiting SNP arrays: a systematic review on the tools to extract underlying genomic structure. Brief. Bioinform., 23, bbac043.

Basu, U. and Parida, S.K. (2021) Restructuring plant types for developing tailor-made crops. Plant Biotechnol. J., 21, 1106–1122.

Boyle, E.A., Li, Y.I. and Pritchard, J.K. (2017) An Expanded View of Complex Traits: From Polygenic to Omnigenic. Cell, 169, 1177–1186.

Chen, S. (2023) Ultrafast one-pass FASTQ data preprocessing, quality control, and deduplication using fastp. iMeta, 2, e107.

Chen, Z.L., Meng, J.M., Cao, Y., et al. (2019) A high-speed search engine pLink 2 with systematic evaluation for proteome-scale identification of cross-linked peptides. Nat. Commun., 10, 3404.

Christersson, L., Sennerby-Forsse, L. and Zsuffa, L. (1993) The role and significance of woody biomass plantations in Swedish agriculture. For. Chron., 69, 687–693.

Dai, X., Hu, Q., Cai, Q., et al. (2014) The willow genome and divergent evolution from poplar after the common genome duplication. Cell Res., 24, 1274–1277.

Dalma-Weiszhausz, D.D., Warrington, J.A., Tanimoto, E.Y. and Miyada, C.G. (2006) The affymetrix GeneChip platform: an overview. Methods Enzymol., 410, 3–28.

Danecek, P., Bonfield, J.K., Liddle, J., et al. (2020) Twelve years of SAMtools and BCFtools. GigaScience, 10, giab008.

Elowson, S. (1999) Willow as a vegetation filter for cleaning of polluted drainage water from agricultural land. Biomass Bioenergy, 16, 281–290.

Freudl, R. (2018) Signal peptides for recombinant protein secretion in bacterial expression systems. Microb. Cell Factories, 17, 1–10.

Guo, Z., Yang, Q., Huang, F., et al. (2021) Development of high-resolution multiple-SNP arrays for genetic analyses and molecular breeding through genotyping by target sequencing and liquid chip. Plant Commun., 2, 100230.

Johnson, M.G., Pokorny, L., Dodsworth, S., et al. (2018) A Universal Probe Set for Targeted Sequencing of 353 Nuclear Genes from Any Flowering Plant Designed Using k-Medoids Clustering. Syst. Biol., 68, 594–606.

Kim, K.-W., Nawade, B., Nam, J., Chu, S.-H., Ha, J. and Park, Y.-J. (2022) Development of an inclusive 580K SNP array and its application for genomic selection and genome-wide association studies in rice. Front. Plant Sci., 13, 1036177.

Kuzovkina, Y.A. and Quigley, M.F. (2005) Willows beyond wetlands: Uses of salix L. species for environmental projects. Water. Air. Soil Pollut., 162, 183–204.

Lee, C.Y., Cheon, K.-S., Shin, Y., et al. (2022) Development and Application of a Target Capture Sequencing SNP-Genotyping Platform in Rice. Genes, 13, 794.

Li, H. (2013) Aligning sequence reads, clone sequences and assembly contigs with BWA-MEM. Arxiv. Available at: http://arxiv.org/abs/1303.3997.

Li, H., Handsaker, B., Wysoker, A., Fennell, T., Ruan, J., Homer, N., Marth, G., Abecasis, G. and Durbin, R. (2009) The Sequence Alignment/Map format and SAMtools. Bioinformatics, 25, 2078–2079.

Masclaux, F.G., Galaud, J.P. and Pont-Lezica, R. (2005) The riddle of the plant vacuolar sorting receptors. Protoplasma, 226, 103–108.

Mathieson, I. (2021) The omnigenic model and polygenic prediction of complex traits. Am. J. Hum. Genet., 108, 1558–1563.

Newsholme, C. (2003) Willows: The Genus Salix, Timber Press. Available at: https://books.google.com/books?id=mcEwgY-AmLAC.

Tarasov, A., Vilella, A.J., Cuppen, E., Nijman, I.J. and Prins, P. (2015) Sambamba: fast processing of NGS alignment formats. Bioinformatics, 31, 2032–2034.

Thomson, M.J. (2014) High-Throughput SNP Genotyping to Accelerate Crop Improvement. Plant Breed. Biotechnol., 2, 195–212.

Varshney, R.K., Bohra, A., Yu, J., Graner, A., Zhang, Q. and Sorrells, M.E. (2021) Designing Future Crops: Genomics-Assisted Breeding Comes of Age. Trends Plant Sci., 26, 631–649.

Wang, N., Li, H. and Huang, S. (2025) Rational Redomestication for Future Agriculture. Annu. Rev. Plant Biol., 76. Available at: 10.1146/annurev-arplant-083123-064726.

Wei, S., Yang, Y. and Yin, T. (2020) The chromosome-scale assembly of the willow genome provides insight into Salicaceae genome evolution. Hortic. Res., 7, 45.

Wu, T., Hu, E., Xu, S., et al. (2021) clusterProfiler 4.0: A universal enrichment tool for interpreting omics data. The Innovation, 2, 100141.

Yang, J., Lee, S.H., Goddard, M.E. and Visscher, P.M. (2011) GCTA: A tool for genome-wide complex trait analysis. Am. J. Hum. Genet., 88, 76–82.

Zhou, X. and Stephens, M. (2012) Genome-wide efficient mixed-model analysis for association studies. Nat. Genet., 44, 821–824.

